# Neural correlates of drinking reduction during cognitive behavioral therapy for alcohol use disorder

**DOI:** 10.1101/2023.02.08.527703

**Authors:** Nasir H. Naqvi, A. Benjamin Srivastava, Juan Sanchez-Peña, Jessica Lee, John J. Mariani, Gaurav H. Patel, Frances R. Levin

## Abstract

Cognitive behavioral therapy (CBT) is an effective treatment for alcohol use disorder (AUD). We hypothesized that the dorsolateral prefrontal cortex (DLPFC), a brain region implicated in cognitive control and goal-directed behavior, plays a role behavior change during CBT by facilitating regulation of craving. To examine this, treatment-seeking participants with AUD (N=22) underwent functional MRI scanning both before and after a 12-week single-arm trial of CBT, using a regulation of craving (ROC) fMRI task designed to measure an individual’s ability to control alcohol craving and previously shown to engage the DLPFC. We found that both the number of heavy drinking days (NHDD, the primary clinical outcome) and the self-reported alcohol craving measured during the ROC paradigm were significantly reduced from pre- to post-CBT [NHDD: t=15.69, p<0.0001; alcohol craving: (F(1,21)=16.16; p=0.0006)]. Contrary to our hypothesis, there was no change in regulation effects on self-reported craving over time (F(1,21)=0.072; p=0.79), nor was there was a significant change in regulation effects over time on activity in any parcel. Searching the whole brain for neural correlates of reductions in drinking and craving after CBT, we found a significant 3-way interaction between the effects of cue-induced alcohol craving, cue-induced brain activity and timepoint of assessment (pre- or post-CBT) on NHDD in a parcel corresponding to area 46 of the right DLPFC (ß=-0.37, p=0.046, FDR corrected). Follow-up analyses showed that reductions in cue-induced alcohol craving from pre- to post-CBT were linearly related to reductions in alcohol cue-induced activity in area 46 only among participants who ceased heavy drinking during CBT (r=0.81, p=0.005) but not among those who continued to drink heavily (r=0.28, p=0.38). These results are consistent with a model in which CBT impacts heavy drinking by increasing the engagement of the DLPFC during cue-induced craving.

## Introduction

Alcohol use disorder (AUD) is a chronic illness associated with significant morbidity and mortality that has substantially increased since 2020 [1–4]. Cognitive Behavioral Therapy (CBT) is an effective treatment for AUD that involves learning strategies for managing or coping with alcohol cravings and a variety of negative emotional states that promote heavy drinking [5– 8]. Prior research on mechanisms of behavior change in CBT has focused on understanding the role of various psychological constructs, e.g., self-regulation, self-efficacy, and acquisition of coping skills. However, attempts at studying these psychological mechanisms have yielded inconsistent results [9,10]. These inconsistencies may result from the inherent complexity in these constructs that may engage multiple underlying cognitive functions. Also, behavior change may occur through processes that occur outside of conscious awareness and thus are inaccessible by self-report alone. Further, these constructs may not directly relate to affective, cognitive, and motivational process implicated in the pathophysiology of AUD [11]. Studying neural mechanisms of behavior change in CBT may help overcome these limitations, allowing for the identification of specific brain systems that can be more specifically targeted to improve existing treatments and develop new, more effective treatments [11].

From a cognitive process perspective, constructs such as self-regulation, self-efficacy, and coping can all be thought of as engaging cognitive control. Cognitive control involves the coordinated deployment of attention, goal representation, action selection, and suppression of competing behaviors, all in the service of reducing automaticity and increasing flexible, goal-directed behavior [12]. Cognitive control can thus be seen as a general function that subserves behavior change, not only in the context of treatments such as CBT, but also among non-treatment seeking individuals who are engaged in the process of reducing heavy drinking. The dorsolateral prefrontal cortex (DLPFC) is a large and heterogenous region of prefrontal cortex that, broadly speaking, is thought to play a central role in cognitive control [12–14]. Both DLPFC structure/function [15–19] as well as behavioral measures of cognitive control [20,21] have been shown to be negatively impacted in AUD. By facilitating cognitive control, CBT may remediate deficits in DLPFC function, increasing the capacity to regulate or reduce alcohol-seeking motivational states such as craving. CBT may also change behavior by promoting a more goal-directed mode of alcohol seeking motivation, as compared to a more automatic mode of alcohol seeking that predominates before treatment. We have previously proposed [22] that this more goal-directed mode is dependent upon the function of the prefrontal cortex broadly, including the DLPFC as well as the ventromedial prefrontal and anterior cingulate cortices, along with the insula. This system becomes engaged under states of high subjective riskiness of drinking and increased value of alternative goals, both of which are promoted by CBT. In the goal-directed mode heavy drinking is more deliberative, more strongly driven by cravings, and more subject to regulation, compared to the automatic mode, where it is non-deliberative and driven largely by non-conscious motivational processes and is less subject to regulation.

While theoretical accounts highlight a potential role for the DLPFC in behavior change in CBT for AUD, research evidence for the impact of CBT on DLPFC functions is mixed. An early study by Schneider et al. [23] examined how CBT impacted neural correlates of cue-induced craving, finding that treatment reduced cue-induced activity in the amygdala and hippocampus, while increasing activity in the superior temporal sulcus. There was no change in activity in the DLPFC. This study included pharmacological interventions, did not explicitly test cognitive control or regulation processes, and did not include clinical outcomes [27]. DeVito et al. [24] examined changes in brain activity during a Stroop task (a cognitive control task [12]) in individuals with a variety of substance use disorders, including AUD, over a course of CBT. They found that CBT decreased Stroop-related activity in the DLPFC, as well as in the anterior cingulate cortex and midbrain. However, this study was not focused on AUD, and did not include clinical outcomes. In another study focused on participants with cocaine use disorder, DeVito et al. [25] showed that changes in Stroop-related activity in DLPFC were associated with number of CBT sessions completed. We recently found that increased resting state functional connectivity (RSFC) between area 9/46 in the DLPFC and the anterior insula was associated with a reduction in heavy drinking during CBT in participants with alcohol use disorder [26]. However, RSFC does not directly measure brain activity that is related to specific mental processes, such a craving or cognitive control [27,28]. In aggregate, these prior studies support a role for the DLPFC in behavior change during CBT for AUD. However, none tested mechanisms of cognitive control processes as they specifically relate to alcohol-seeking motivational states, e.g., craving, which may be especially useful probes for understanding mechanisms of CBT.

Cognitive control may play a particular role in CBT through the regulation of cue-induced alcohol craving. One of the core skills taught in CBT is the ability to down-modulate or regulate craving in high-risk situations (cue-induced craving) using various cognitive strategies, including by thinking about the long-term negative consequences of drinking [29]. This is a form of reappraisal-based emotion regulation, which is theorized to depend upon cognitive control [30]. Kober and colleagues adapted this therapeutic strategy in a regulation of craving fMRI paradigm, applying it across a variety of substances, including alcohol [21,31–34]. Regulation of craving is associated with increased activity in areas within DLPFC, anterior cingulate and ventrolateral prefrontal cortices, coupled with decreased activity in the ventral striatum. We have previously shown this form of regulation to be impaired in individuals with AUD [21]. Altogether, this suggests that the regulation of craving task may be a particularly sensitive probe for revealing the role of the DLPFC in behavior change during CBT. However, to date, this task has never been combined with a clinical trial of CBT.

In this study, we used fMRI to examine how brain function related to cognitive regulation of cue-induced craving changes over the course of CBT for AUD and how these changes are related to clinical drinking outcomes. We recruited treatment-seeking participants with AUD who were drinking heavily at baseline. The participants then completed the regulation of craving task both before and after undergoing approximately 12 weeks of manualized CBT for AUD. In this task, participants are shown alcohol cues that are known to elicit cravings while being instructed to focus on long-term negative consequences of drinking vs. immediate pleasurable consequences, using food cues as control stimuli, while brain activation and craving self-report data are collected. We chose reduction in heavy drinking as the primary clinical outcome since it is a clinically meaningful outcome [35,36] and is more attainable than total abstinence. We hypothesized that a) CBT will increase the ability to regulate cue-induced alcohol craving; b) this improved ability to regulate craving during CBT will be associated with an increase in DLPFC activity; and c) increased DLPFC activity will in turn be related to reductions in heavy drinking.

## Methods

### Participants

Details of participant recruitment, including inclusion/exclusion criteria, study structure, data collection, and CBT are published elsewhere [26]. Briefly, adults seeking treatment for alcohol-related problems were recruited from the New York City Area. All participants met Diagnostic and Statistical Manual for Mental Disorders, Fifth Edition, criteria for AUD, as confirmed by the MINI. Participants were excluded for neurological or medical illnesses that would interfere with MRI scanning, other moderate or severe substance use disorders besides nicotine or caffeine, significant psychiatric illness, or a significant alcohol withdrawal history. The study was approved by the New York State Psychiatric Institute (NYSPI) Institutional Review Board. All participants provided informed consent.

### Clinical outcome and analysis

The primary clinical outcome was the number of heavy drinking days (NHDD), defined as the number of days over the previous 28 days on which participants consumed ≥4 standard drinks/day for women and ≥5 drinks/day for men, as assessed using the 28 Day TLFB. NHDD was calculated for both pre-CBT and post-CBT timepoints, measured at the pre-CBT and post-CBT behavioral assessments, respectively and was contrasted using a paired t-test.

### Regulation of craving (ROC) task and analysis

At both the pre- and post-CBT timepoints each participant performed the ROC task while fMRI data were collected. The task was programmed in E-Prime version 2.0 (Psychology Software Tools, Sharpsburg, PA) and displayed to the participants using a back projection mirror. Prior to performing the ROC task in the scanner, participants completed 8 practice trials outside the scanner. The structure of the ROC task is as follows **(Figure S1**): Participants were shown an instructional word for 2 seconds, directing them to focus on either the immediate, pleasurable consequences of consuming the depicted item (“LOOK”) or the long-term, negative consequences (“NEGATIVE”) of repeatedly consuming the depicted item. They were then immediately shown a picture cue for 6 seconds: either an image of an alcohol or an image of high-calorie food, followed by a fixation cross during an inter-stimulus interval randomly jittered between 1 and 7.5s. They were then shown a visual instruction to rate the desire to consume the depicted item (craving) on a 1-5 Likert scale, during which they made rating responses with their right hand using a five-finger button-response unit. They were given up to 3 seconds to make their response. Once the response was made (or after 3 seconds, which ever came first), participants were then shown a fixation point during the inter-trial interval, which was randomly jittered between 1.5 and 7s; this was then followed by the next trial. The cue images (alcohol and food) were previously validated as eliciting moderate cravings in participants with AUD [21]. The order of instructions was counterbalanced across alcohol (n=50) and food (n=50) images. Participants performed 4 runs of the task with 20 trials per run. We then examined how response data (craving scores) varied as a function of cue (food vs alcohol cues), instruction (LOOK vs. NEGATIVE), and measurement timepoint (pre-CBT vs. post-CBT) with a repeated measures ANOVA. Post hoc comparisons for this and subsequent ANOVAs were assessed using paired t-tests with a threshold of p < 0.05.

### MRI acquisition and preprocessing

The scanning procedures for both pre- and post-treatment MRI scanning sessions were the same. Participants first completed a 7-day TLFB [37] to quantify recent drinking and reported the time since the last drink. They were then administered a breath alcohol test, vital signs, the CIWA-Ar, the Alcohol Urge Questionnaire [38], and a urine drug screen. This was followed by the ROC practice trials outside the scanner, followed by fMRI scanning during ROC task during, followed by a 10-minute resting state scan, the results of which are published elsewhere [26,39].

MRI data were collected on a 3T MR750 GE Scanner. Functional images were acquired with a single-band EPI sequence with a TR of 2000 ms, TE of 25 ms, flip angle of 77°, 64 × 64 in-plane matrix, field of view 19.2 cm, and 45.3 mm slices in an ascending, interleaved order. High-resolution (1 mm isotropic) structural T1 images were also acquired with a flip angle of 12°, 256 × 256 in-plane matrix, and field of view 25.6 cm. Data were processed using fMRIPrep version 1.5.10, which performs standard preprocessing steps including alignment to individual’s anatomical data, movement correction, distortion correction, and atlas alignment into MNI volume space [40]. We then used Ciftify version 2.3.3, which allows for the adaptation of Human Connectome Project Pipelines to legacy data in which T2w anatomical images and field maps were not acquired [41]. Specifically, Ciftify performs surface-based extraction and surface atlas alignment of gray matter voxels to improve co-registration of functional maps between individuals and with standard surface atlases ([41,42]).

### ROI selections for parcel-based analyses

We used a surface-based parcel-wise analysis, which has been shown to improve cross-subject alignment and to provide more reliable anatomical specificity of brain function than voxel-based approaches [43,44]. Cortical areas were defined using a multimodal parcellation from Glasser et al. [42] Subcortical parcellations were defined using separate atlases containing dorsal and ventral striatal structures [45], amygdala [46], and bed nucleus of the stria terminalis (BNST) [47].

### Analysis of brain activation during the ROC task

A parcel-wise general linear model (GLM) was used to estimate the BOLD response at each parcel for each cue-instruction event pair (LOOK/alcohol, NEGATIVE/alcohol, LOOK/food, NEGATIVE/food), at each timepoint, for each participant. Following previous studies using the ROC task [31,33], the instruction and picture cue events were combined into a single 8 second event. The self-report response was modeled separately. Each event type was modeled as a separate boxcar regressor that was then convolved with the hemodynamic response function [48]. These regressors were entered into the first level GLM analyses as predictors of the measured BOLD response, resulting in parameter estimates (β-weight weights) for each of the 4 cue-instruction event conditions, at each parcel, in each participant, at each timepoint. For each brain parcel, we then examined how cue-instruction event-evoked brain activity varied as a function of cue (food vs alcohol cues), instruction (LOOK vs NEGATIVE), and measurement timepoint (pre-CBT vs post-CBT) using a repeated measures ANOVA. A false discovery rate (FDR) correction (adjusted p=0.05) using the Benjamini-Hochberg procedure [49] was performed for each factor to correct for the multiple comparisons across the 396 parcels. To allow for more direct comparison between our results and those of Kober et al. [31], we also performed a whole-brain voxel-wise analysis using paired t-tests, contrasting the NEGATIVE and LOOK conditions (NEGATIVE-LOOK), combining across alcohol and food trials (see **Supplement** for details and results).

### Analysis of the relationships between changes in brain activity, changes in craving and changes in heavy drinking, from pre-CBT to post-CBT timepoints

For each brain parcel, we used a linear mixed effects model to examine how NHDD varied as a function of cue-induced alcohol craving, alcohol cue-induced brain activity, and pre- vs. post-CBT timepoint. Following convention in prior cue-reactivity imaging studies [50–52], we defined alcohol cue-induced brain activity as the average BOLD response in that parcel (β- weight) during alcohol trials minus the average BOLD response during non-alcohol (food) trials. Cue-induced alcohol craving was defined as the average Likert scale rating for alcohol cues only, not subtracting craving for alcohol cues [53,54]. For both the brain activation measure of alcohol cue-induced brain activity and the behavioral measure of cue-induced alcohol craving, trials were averaged without regard to LOOK vs. NEGATIVE instruction, i.e., regulation was not included as a factor in this regression analysis. Time between scans was also entered as an independent variable. Continuous variables were z-scored. The individual participants were modeled as a random effect. The results for each factor were false discovery rate (FDR) corrected (adjusted p < 0.05) for the multiple comparisons across the 396 brain parcels. Finally, we performed three pairs of follow-up regression analyses, restricted to parcels that showed a 3-way interaction between cue-induced craving, cue-induced brain activity and CBT timepoint on NHDD. The first pair of regressions examined the relationship between alcohol cue-induced activity brain activity and NHDD, separately for pre- and post-CBT timepoints. The second pair examined the relationship between cue-induced alcohol craving and NHDD, separately for pre- and post-CBT timepoints. The third pair examined reductions in cue-induced alcohol craving from pre- to post-CBT and reductions in alcohol cue-induced brain activity from pre- to post-CBT, first in participants who continued heavy drinking and then in participants who ceased heavy drinking. Alpha levels for these follow-up regressions were Bonferroni corrected for 6 tests (α = 0.05/6 = 0.008).

## Results

### Demographics

#### Demographics

**Table 1** describes demographic characteristics of the 22 participants who were included in the analyses. For detailed demographic data and the CONSORT diagram, see **Supplement. Heavy drinking outcomes and self-reported craving in the ROC task**

**Table 1:**
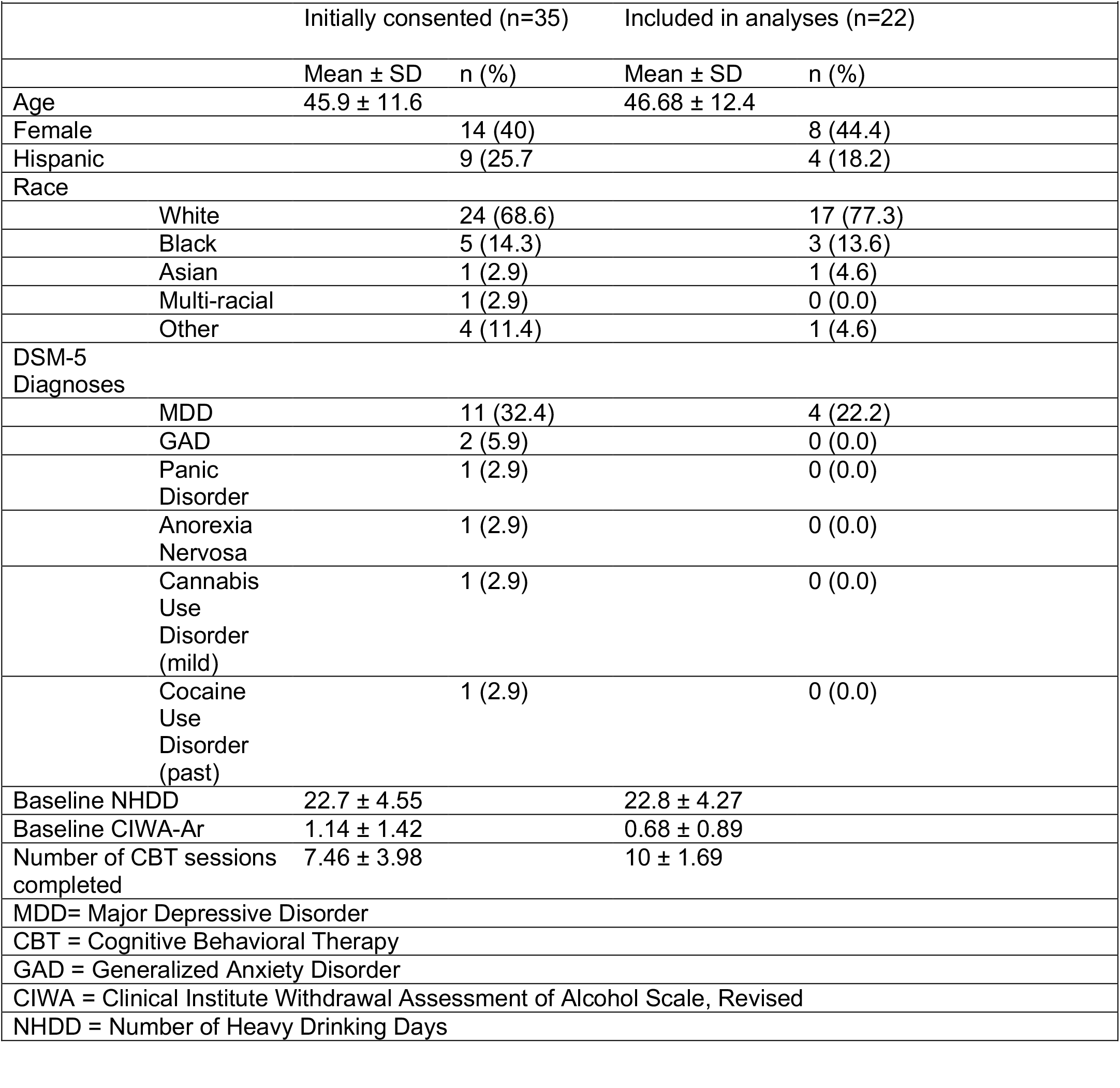
Subject Characteristics

Number of heavy drinking days (NHDD) was significantly reduced over the course of CBT (t(21)=15.69; p<0.0001). We found significant main effects of regulation instruction (F(1,21)=7.59; p=0.012) and time (F(1,21)=13.40; p=0.0015) as well as a significant interaction effect of cue type and time (F(1,21)=16.16; p=0.0006). We did not find a significant main effect of cue, interaction effects of instruction and cue, interaction effects of instruction and time, or significant 3-way interaction effects of instruction, cue, and time. Post-hoc testing showed that the mean craving scores were 1) significantly greater in LOOK than in NEGATIVE conditions (t(87)=4.86; p < 0.0001), 2) significantly greater at the pre-CBT timepoint than post-CBT timepoint (t(87)=5.35; p < 0.0001), and 3) significantly lower for alcohol cues than for food cues, but only at the post-CBT timepoint, as compared with the pre-CBT timepoint (t(43)=6.80; p < 0.0001). There was no significant change in craving score for food cues from pre-CBT to post-CBT timepoints (t(43)=1.23; p = 0.22) (**Figure 1**).

**Fig 1.**
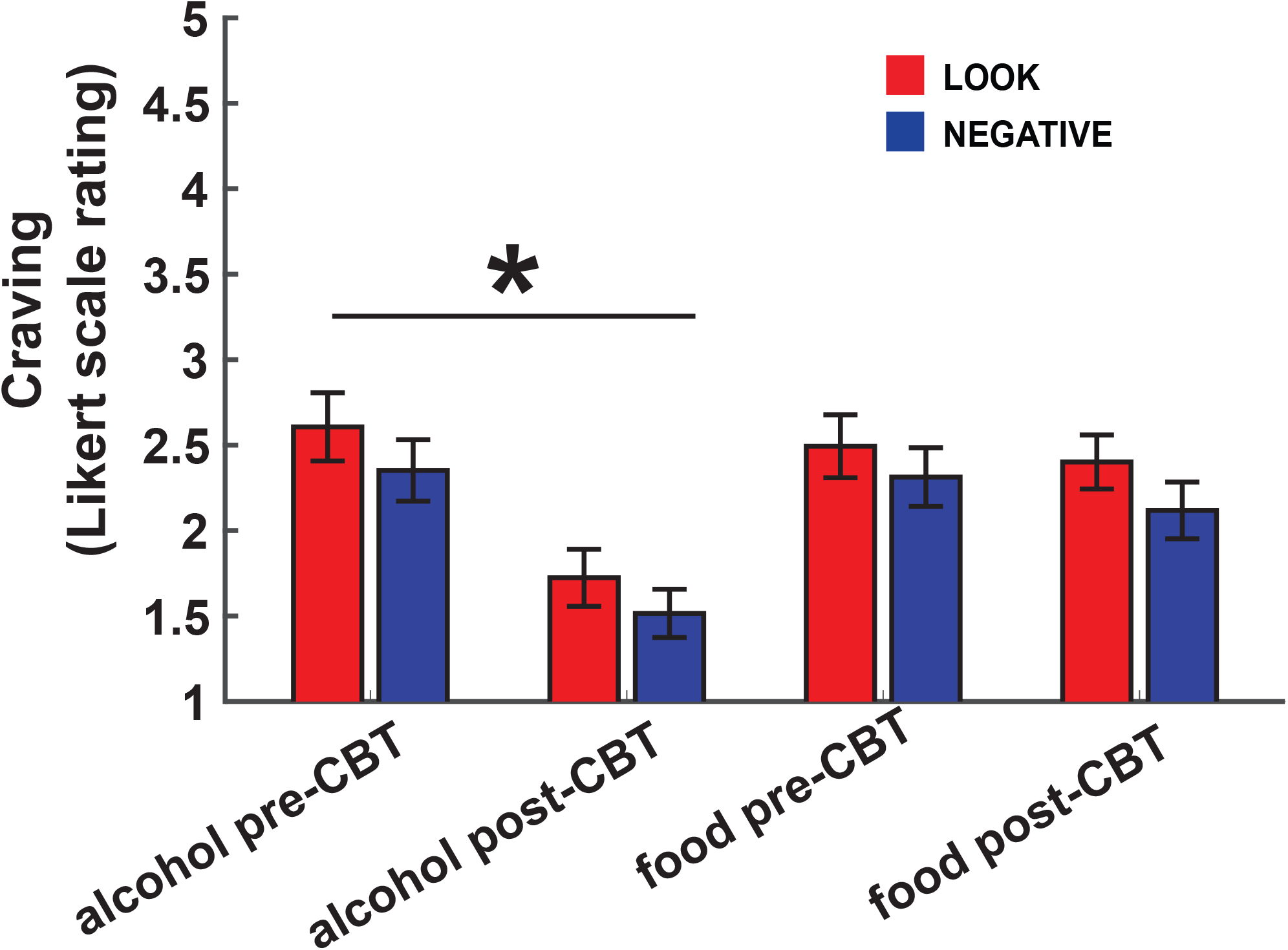
Mean craving (Likert scale ratings) reported by participants as a function of cue type (alcohol/FOOD), instruction type (LOOK/NEGATIVE). Overall, significant main effects were present for instruction and time, and a significant interaction effect between cue type and time was found. Post hoc testing using Tukey’s HSD showed that craving for alcohol cues was significantly lower at the post-CBT time point when compared to the pre-CBT timepoint. Significance is noted for the cue x time interaction at *p<0.0001. Error bars represent 95% confidence intervals.

### Brain activation during the ROC task

There was a significant main effect of cue-type in multiple parcels on both sides within posterior occipital, temporal and parietal lobes, corresponding to primary and association visual cortical areas, as well as within a region of the right mid-posterior insula (PoI2) (**Figure 2, Table S1**). We found a significant main effect of regulation instruction on brain activity within a single parcel corresponding to the left ventral superior temporal sulcus (STSvp) (F(1,21)=22.18; p=0.047, FDR corrected). Post-hoc testing showed that activation in this parcel was greater in the NEGATIVE than in the LOOK condition (t(87)=-5.45; p< 0.0001). Voxelwise analyses demonstrated similar foci of activation as was described in Kober et al. [31] (**Figure S3**). However, there were no differences in these regulation strategy contrasts pre-versus post-CBT in the DLPFC or anywhere else. In addition, there were no main effects of cue-type across the brain.

**Fig 2.**
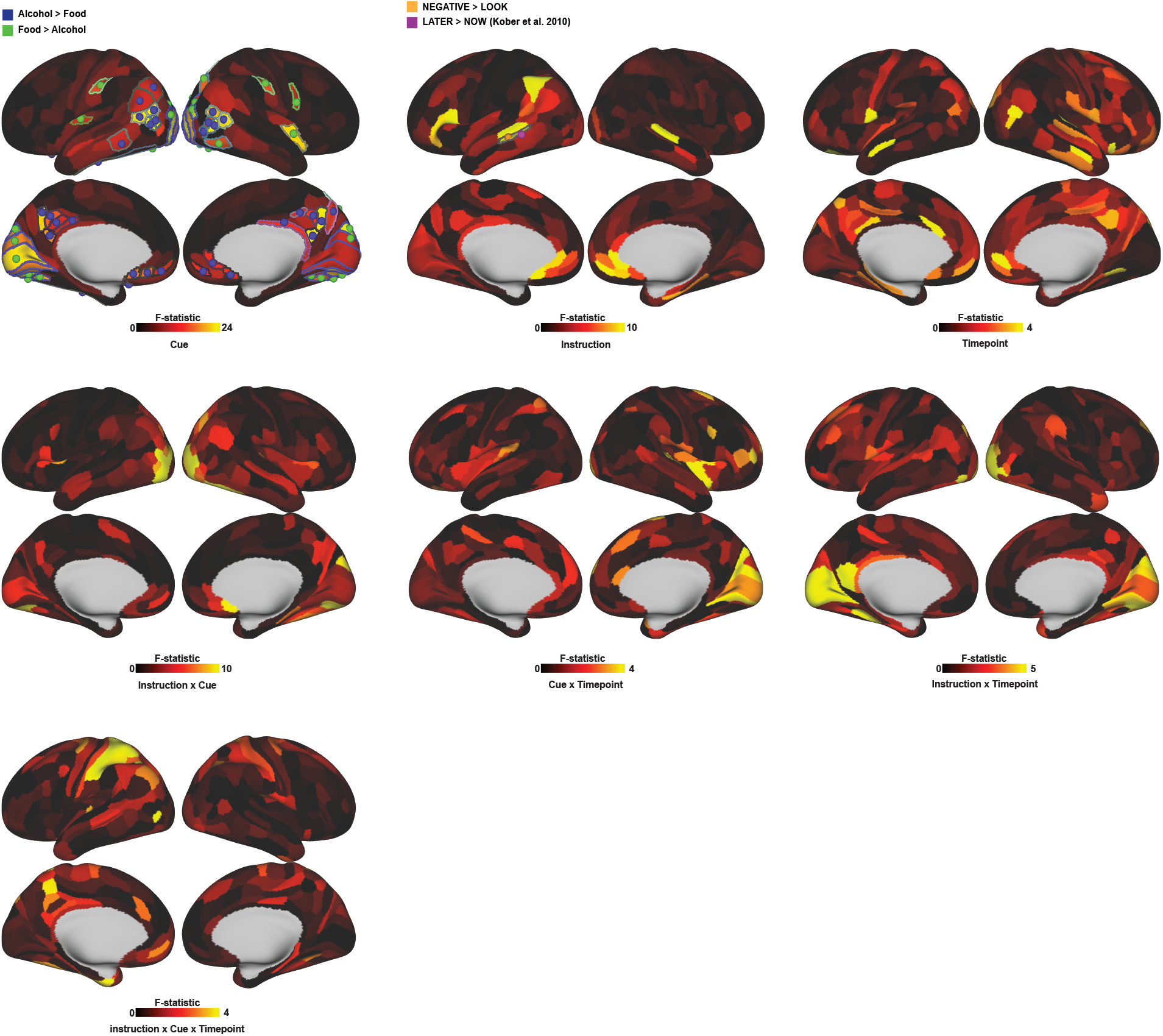
Main and interaction effects of cue type, instruction, and pre-CBT/post-CBT timepoint on whole brain activity. Overall, significant main effects of cue type were found in parcels within the posterior occipital, temporal, and parietal lobes. Main effects of instruction were found in a single parcel in the left superior temporal sulcus (STS). Post hoc Tukey’s HSD test revealed that areas where BOLD response to alcohol cues was greater than food cues in parietal and temporal areas (blue dots), whereas the BOLD response was greater for food than alcohol cues in the occipital lobe (green dot). Significant parcels are outlined. For instruction, BOLD response in the STS was greater for the NEGATIVE compared with the LOOK condition (orange dot). For comparison, the peak STS activation in the LATER vs NOW condition from Kober et al 2010 is denoted by the pink dot. See the **supplement** for parcel labels and detailed statistics.

### Neural correlates of cue reactivity

When we examined how NHDD varied as a function of cue-induced brain activity, cue-induced alcohol craving, and timepoint (pre-CBT/post-CBT) and their interactions, we found a significant 3-way interaction effect of cue-induced alcohol craving, pre-/post CBT timepoint, and cue-induced brain activity in a parcel corresponding to area 46 of the right DLPFC (β = -0.37, 95% CI -0.57 - -0.13, p = 0.0006 uncorrected, 0.046 corrected), as well as small number of parcels within occipital, temporal and parietal regions, corresponding to higher-order visual cortices (**Figure 3**; see **Supplement** for detailed statistics for the 3-way interactions in regions outside of DLPFC). When we examined how NHDD varied as a function of cue-induced brain activity, cue-induced alcohol craving, and timepoint (pre-CBT/post-CBT) and their interactions, we found a significant 3-way interaction effect of cue-induced alcohol craving, pre-/post CBT timepoint, and cue-induced brain activity in a parcel corresponding to area 46 of the right DLPFC (β = -0.37, 95% CI -0.57 - -0.13, p = 0.0006 uncorrected, 0.046 corrected), as well as small number of parcels within occipital, temporal and parietal regions, corresponding to higher-order visual cortices (**Figure 3**; see **Supplement**) for detailed statistics for the 3-way interactions in regions outside of DLPFC). Within the right DLPFC area 46 parcel, we found a significant 2-way interaction between cue-induced alcohol craving and pre- vs. post-CBT timepoint (β = 0.38, 95% CI 0.18 - -0.59, p = 0.0006 uncorrected, 0.0015 corrected), and a trend-level 2-way interaction between the effects of cue-induced alcohol craving and cue-induced brain activity on NHDD (β = 0.24, 95% CI 0.092 - -0.40, p = 0.0025 uncorrected, 0.073 corrected). We also found trend-level main effects of cue-induced brain activity (β = -0.34, 95% CI -0.56 - -0.12, p = 0.0039 uncorrected, 0.11 corrected) and time between scans (β = -0.50, 95% CI -1.00 - -0.0023, p = 0.049 uncorrected, 0.088 corrected). There were no significant or trend-level main effects of either cue-induced alcohol craving or timepoint on NHDD in this parcel. We found no significant 2-way interaction between the effects of cue-induced activity and timepoint on NHDD in this parcel.

**Fig 3.**
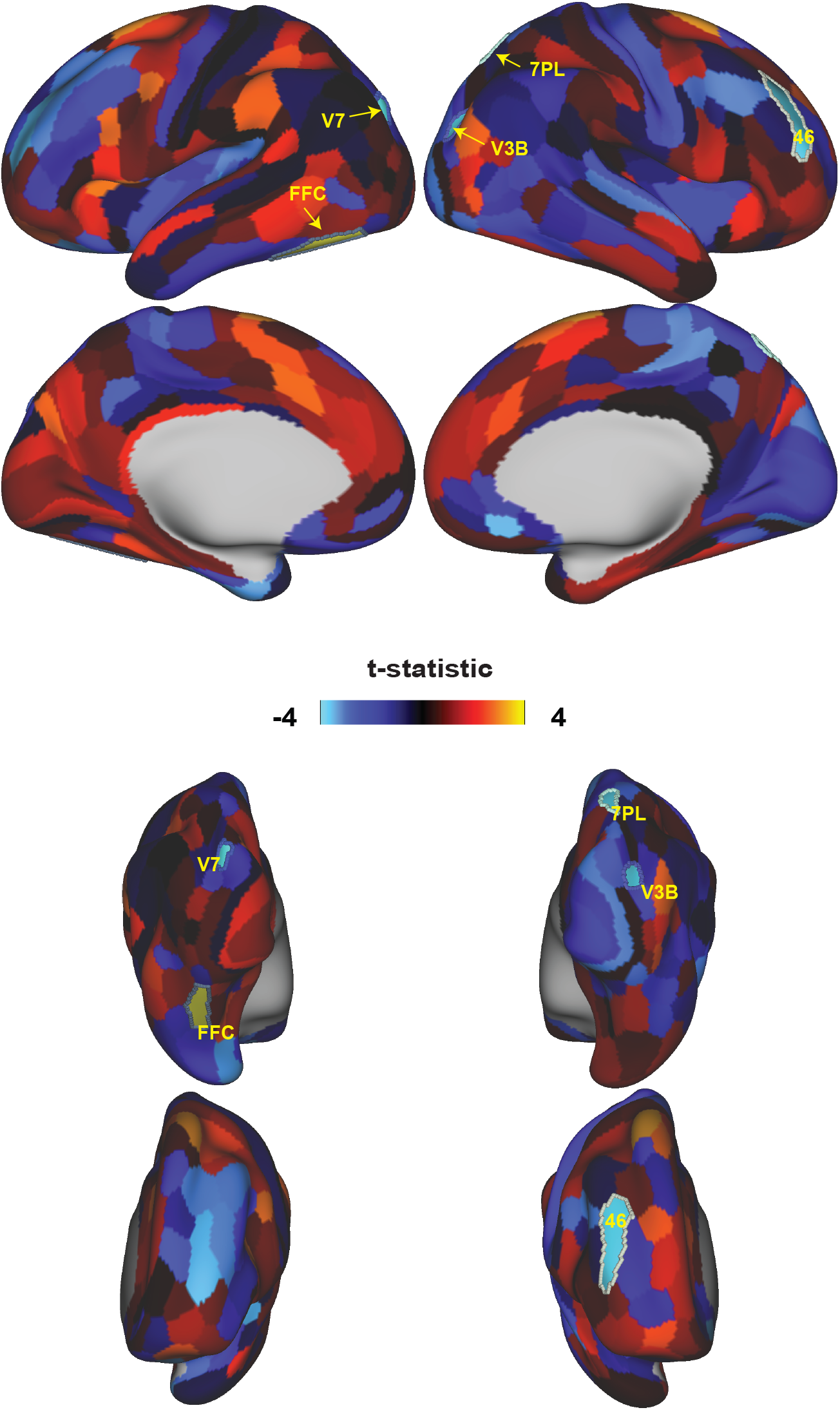
Parcellated map of 3-way interaction (cue induced alcohol craving x alcohol cue-induced brain activity x pre-CBT/post-CBT timepoint) on NHDD. Shown are lateral (top row) and medial (second row) views as well as posterior (3rd row) and anterior (4^th^ row) Significant parcels in this 3-way interaction are outlined and parcels corresponding to the right DLPFC area 46 (R_46; β = -0.37, 95% CI -0.57 - -0.13, p = 0.0006 uncorrected, 0.046 corrected) as well as right (R_V3B, R_7PM) and left (L_V7, L_FFC) higher order visual areas. See **supplement** for detailed statistics of areas outside the DLPFC.

### Follow-up analyses of the role of right area 46 in CBT related changes in craving, cue reactivity, and drinking

All subsequent analyses were focused on area 46 of the right DLPFC, since this was the only area to show a significant 3-way interaction between cue-induced craving, cue-induced brain activity and timepoint on NHDD. When we examined the relationship between cue-induced alcohol craving and NHDD at both pre- and post-CBT timepoints, there was no significant (p < 0.0125, corrected) or trend level (p < 0.05, uncorrected) relationship between cue-induced alcohol craving and NHDD at the pre-CBT (r = -0.37, p = 0.092) timepoint; however, at the post-CBT timepoint there was a trend-level association between cue-induced alcohol craving and NHDD (r = 0.42, p = 0.049). **(Figure 4A)**. When we examined the relationship between cue-induced DLPFC activity and NHDD at the pre- and post-CBT timepoints, there was no relationship between cue-induced DLPFC area 46 activity and NHDD at either the pre-CBT (r = -0.28, p = 0.22) or post-CBT (r = -0.034, p = 0.88) **(Figure 4B)**.

**Fig 4.**
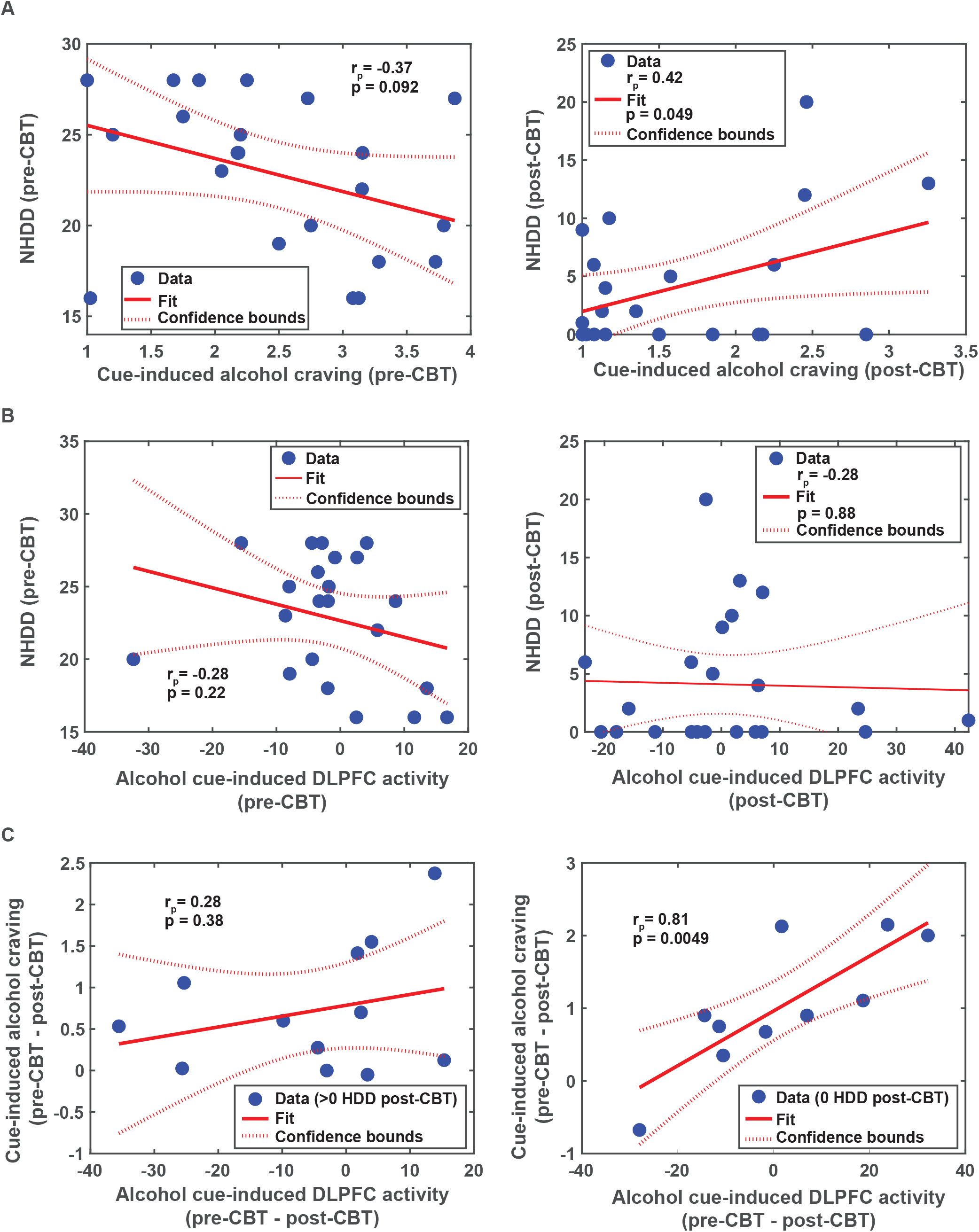
Follow-regression analyses focused on area 46 of the right DLPFC. Shown are the regression lines for 1) the relationships between cue-induced alcohol craving and NHDD at pre-CBT and post-CBT timepoints (**A**), the relationships between alcohol-cue induced DLPFC activation at pre-CBT and post-CBT timepoints (**B)**, and the relationship between the change (pre-CBT – post-CBT) in alcohol-cue induced DLPFC activation in participants with >0 HDD post-CBT and with 0 HDD post-CBT (**C**).

We then examined the relationship between changes in cue-induced DLPFC area 46 activity and changes in cue-induced alcohol craving from pre-CBT to post-CBT timepoints. This was done separately for participants who ceased heavy drinking at the post-CBT timepoint and those who continued heavy drinking. For participants with NHDD = 0 at the post-CBT timepoint (ceased heavy drinking), the pre- to post-CBT reduction in cue-induced DLPFC area 46 activity was significantly associated with the pre- to post-CBT reduction in cue-induced alcohol craving (r =0.81, p = 0.0049). In participants with NHDD > 0 at the post-CBT timepoint (continued heavy drinking) the pre- to post-CBT reduction cue-induced DLPFC area 46 activity did not significantly predict the pre- to post-CBT reduction in cue-induced alcohol cue induced craving (r = 0.28, p = 0.38) **(Figure 4C)**.

## Discussion

Here we showed that in participants with AUD, treatment with CBT is associated with 1) significant reductions in drinking, indicative of a clinical effect for the purposes of a mechanistic translational study, 2) an overall decrease in cue-induced alcohol craving as measured in the ROC paradigm, and 3) a strong relationship between reductions in craving and reductions in DLPFC functioning, but only in patients who cease heavy drinking. These results provide evidence that the effectiveness of CBT in AUD is dependent upon DLPFC function and that the DLPFC plays a role in modulating the incentive salience of alcohol cues as indexed by self-reported craving.

Prior studies [31,33] have shown that regulation of cue-induced craving was associated with increased activity in a DLPFC region in the posterior middle frontal gyrus, approximating Brodmann’s area (BA) 8 [55,56], and that activity in this region was related to individual differences in regulation success. While we were able to replicate the finding of regulation related activity in the DLPFC approximating BA 8, we did not find that activity in this area changed with CBT or was associated with a change in heavy drinking. The DLPFC region where we did find a relationship between activity, craving, and reductions in heavy drinking was located in a parcel corresponding to area 46, a region of the DLPFC that has previously been shown to play a core role in multiple forms of cognitive control [57], as well as a role in emotion regulation [58]. Another discrepancy between our results and prior fMRI studies of regulation of craving is that we did not show any significant down-modulation of subcortical regions such as the ventral striatum and amygdala during regulation of craving, as has been shown in previous studies of regulation of craving [31,33,59]. The one area where we did find a regulation effect was the left ventral STS, which is consistent with previous studies showing STS activity to be related to regulation of craving [31] as well as other forms of emotion regulation [60]. The STS is also a region where Schneider et al. [23] found alcohol cue-induced activity to increase over the course of CBT for AUD, suggesting that changes in drinking may have been related to changes in regulation.

The discrepancies between our results and those of prior studies of regulation of craving may be explained by several factors. Previous studies revealed DLPFC activity that was specifically related to the reduction of craving in response to a controlled regulation strategy (“think about long-term negative consequences”). In contrast, we found that DLPFC activity was related to reductions in craving over time, but these findings were not attributable to a controlled regulation strategy. These reductions in craving may have come about through other forms of regulation that are relatively non-controlled/implicit but were nevertheless the result of treatment. Further, prior studies were all done in non-treatment seeking participants [21,34], whereas our study was in treatment-seekers. Treatment seeking participants are likely to be in a more highly regulated/goal-directed state at baseline/pre-treatment, as compared to non-treatment seekers [61]. This high baseline level of regulation in the present study may have served to reduce the overall level of cue-induced alcohol craving at both pre- and post-CBT timepoints and to dampen both the effects of regulation during the ROC task and the effects of CBT on this regulation. Indeed participants in our study rated relatively low levels of overall cue-induced alcohol craving at baseline (before CBT), compared to prior regulation studies, including our own prior behavioral study using the same stimuli and task design involving non-treatment seeking participants [21]. Also, the self-reported cravings in the present study were no greater for alcohol than for food cues, and the regulation effects we found were relatively small, compared to other studies, including our own prior behavioral study [21,31,33]. Future studies should examine differences in regulation of craving between treatment and non-treatment seekers.

CBT involves bringing to awareness the triggers and consequences of craving and heavy drinking, promoting alternative goals, and helping the patient engage in self-regulatory strategies aimed at managing craving, negative emotions, and impulsivity [8,10,62]. We have proposed that treatments such as CBT promote a shift from a more automatic to a more goal-directed mode of alcohol seeking, in which heavy drinking is more subject to deliberation and self-regulation, and which is more strongly tied to subjective feelings, such as craving [22]. This model is supported by our findings that patients who ceased heavy drinking during CBT exhibited a stronger relationship between reductions in craving and reduction in DLPFC activity, compared to patients who continued heavy drinking; patients who cease drinking are presumably in a more goal-directed mode, compared to patients who continue to drink heavily. This model is also supported by the trend-level finding that the relationship between cue-induced alcohol craving and heavy drinking is stronger after CBT than before CBT.

Conclusions from this study should be drawn carefully given its limitations. Our study included neither a non-treatment seeking group nor a control intervention. Thus, we do not know whether the changes in heavy drinking, alcohol craving and associated change,s and DLPFC activation that we observed were due to the specific effects of CBT, non-specific effects of treatment, non-treatment-related behavior change processes, or the mere passage of time. Next, the ROC task was designed to assess regulation of self-reported craving using a controlled regulation strategy that may not resemble the strategies actually used by patients, which may be much more implicit or non-controlled. Further, heavy drinking is driven by more than subjective craving when presented with an alcohol stimulus, and it also involves a number of covert motivational processes, such as attentional bias toward alcohol cues. Finally, conclusions are limited by a small sample size. To address these limitations, future studies should include a larger sample size, a matched non-treatment seeking group and a control intervention, along with an fMRI paradigm with greater ecological validity that addresses more implicit/less controlled forms of regulation and that includes multiple behavioral measures of alcohol-seeking motivation.

## Author Contributions

NHN was responsible for the overall study concept and design, as well as the implementation of the clinical treatment and the collection of the clinical and imaging data. JJM and FRL were involved in the design and implementation of the clinical treatment. ABS and JS processed the data, and ABS analyzed the data, all with supervision from GHP and JL. ABS and NHN wrote the manuscript with input from GHP and FRL. GHP, NHN and FRL are all research mentors of ABS. All authors reviewed content and approved the final version for publication.

