## Supplement for "Neural correlates of drinking reduction during cognitive behavioral therapy for alcohol use disorder"

**Demographic data**

We screened 445 participants, of whom 108 (24.3%) were not seeking treatment or were entered into another treatment trial; 66 (14.8%) did not complete the screening process; 75 (16.9%) did not meet minimum AUD criteria; and 158 (35.5%) were excluded for psychiatric or medical conditions, cognitive impairment, or other exclusionary criteria. Of the 38 participants initially enrolled in the treatment study, three did not undergo scanning (one was medically unstable, one had a tattoo, and one had head too large for the scanner). Of the 35 participants who underwent baseline scanning, 23 underwent both pre- and post-CBT MRI sessions. One participant was discontinued from the study for safety reasons and was excluded from these analyses because of a lack of a post-CBT assessment of drinking days. None of the participants were excluded because of an inability to abstain for more than 24 h or because they demonstrated significant alcohol withdrawal after 24 h of abstinence. For CONSORT diagram for this trial, including reasons for dropout, see **Figure S2**.

**Voxelwise analysis of whole brain differences in instruction**

Though the main results showed similar patterns for the main effect of instruction (**Fig 2.)**, with significant findings in the superior temporal sulcus and trend trend level findings in the inferior frontal gyrus, we did not see main effects of instruction in the DLPFC (broadly, Brodmann’s area 8) as shown by Kober et al. One possibility for this discrepancy is that due to this area, shown in a voxelwise is not adequately captured by the Glasser parcellation. To determine if we were able to spatially replicate the main results from Kober et al 2010 for the NEGATIVE-LOOK (described as LATER-NOW) condition, we performed first and second level analyses as described in the main text methods, only using the unparcellated time series for the GLM. We then used paired t-tests to examine the NEGATIVE and LOOK conditions (NEGATIVE-LOOK). We then transformed the Talairach coordinates of the peak activations listed in Kober et al to MNI space via icbm2tal (brainmap.org). Broadly the peak activations for LATER-NOW corresponding with peak activations from our findings, including the DLPFC finding approximating Brodmann’s area 8. Shown in **Figure** **S3.**

**Detailed analyses of anatomical locations and statistics for parcels significant in the 3-way interaction (cue-induced alcohol craving x cue-induced brain activity x pre-/post-CBT timepoint) on NHDD.**

For Parcel V3B in the right visual cortex, in addition to the significant 3-way interaction effect (β = -0.40; 95% CI -0.62 - -0.19, p = 0.00053 uncorrected, 0.046 corrected) there was a significant interaction effect of cue-induced alcohol craving and timepoint (β = -0.31; 95% CI -0.10 - -0.51, p = 0.0046 uncorrected, 0.0063 corrected) and a trend level main effect of time between scans (β = -0.73; 95% CI -1.26 - -0.19, p = 0.010 uncorrected, 0.072 corrected) on NHDD. There were no significant interaction effects between cue-induced alcohol craving and timepoint). There were no significant main effects of cue-induced alcohol craving, cue-induced brain, or timepoint.

For parcel R_7PL in the right parietal lobe, in addition to the significant 3-way interaction effect (β = -0.53; 95% CI -0.81 - -0.24, p=0.0006 uncorrected, 0.046 corrected), we found a significant interaction effect of cue-induced alcohol craving and timepoint (β = 0.30; 95% CI 0.98 - 0.51, p = 0.0049 uncorrected, 0.0067 corrected). There was a trend level main effect of a trend level interaction effect of cue-induced alcohol craving and cue induced brain activity (β = 0.22; 95% CI 0.021 - 0.40, p = 0.031 uncorrected, 0.0014 corrected) and time between scans (β =- 0.80; 95% CI -1.33 - -0.26, p = 0.0046 uncorrected, 0.071 corrected). We did not find a significant interaction effect between cue-induced brain activity and timepoint. There was no significant main effect of cue-induced alcohol craving, cue-induced brain activity, or timepoint.

For parcel L_V7 in the left visual cortex, in addition to the significant 3-way interaction effect (β = -0.62; 95% CI -0.92 - -0.33, p=0.0001 uncorrected, 0.046 corrected), we found a significant interaction effect of cue-induced alcohol cravings and timepoint (β = -0.62; 95% CI 0.18 - 0.63, p=0.0008 uncorrected, 0.0017 corrected). We found trend level main effects of cue-induced alcohol craving (β = -0.19; 95% CI -0.33 - -0.044, p=0.012 uncorrected, 0.064 corrected) and timepoint (β = -1.25; 95% CI -2.35 - -0.14, p=0.028 uncorrected, 0.53 corrected) on NHDD. We did not find significant interaction effects of cue-induced alcohol craving and cue-induced activity or cue-induced brain activity and timepoint. We did not find significant main effects of cue-induced brain activity or time between scans.

For parcel L_FFC in the fusiform face complex, in addition to the significant 3-way interaction effect (β = 0.72; 95% CI 0.35 – 1.10, p=0.0004 uncorrected, 0.046 corrected), we found significant interaction effects of cue-induced brain activity and timepoint (β = 0.88; 95% CI 0.50 – 1.27, p < 0.0001 uncorrected, 0.0013 corrected) and main effects of cue-induced alcohol craving and timepoint (β = 0.47; 95% CI 0.28 – 0.66, p < 0.0001 uncorrected, 0.0002 corrected). We found trend level main effects of cue-induced alcohol craving (β = -0.17; 95% CI -0.31 – -0.025, p = 0.023, uncorrected, 0.067 corrected) and time between scans (β = -0.73; 95% CI -1.23 – -0.22, p = 0.0061, uncorrected, 0.072 corrected). There was no significant interaction effect of cue-induced alcohol craving and cue-induced brain activity timepoint. We did not find main effects of cue-induced brain or timepoint.

**Figure S1: ROC schematic**

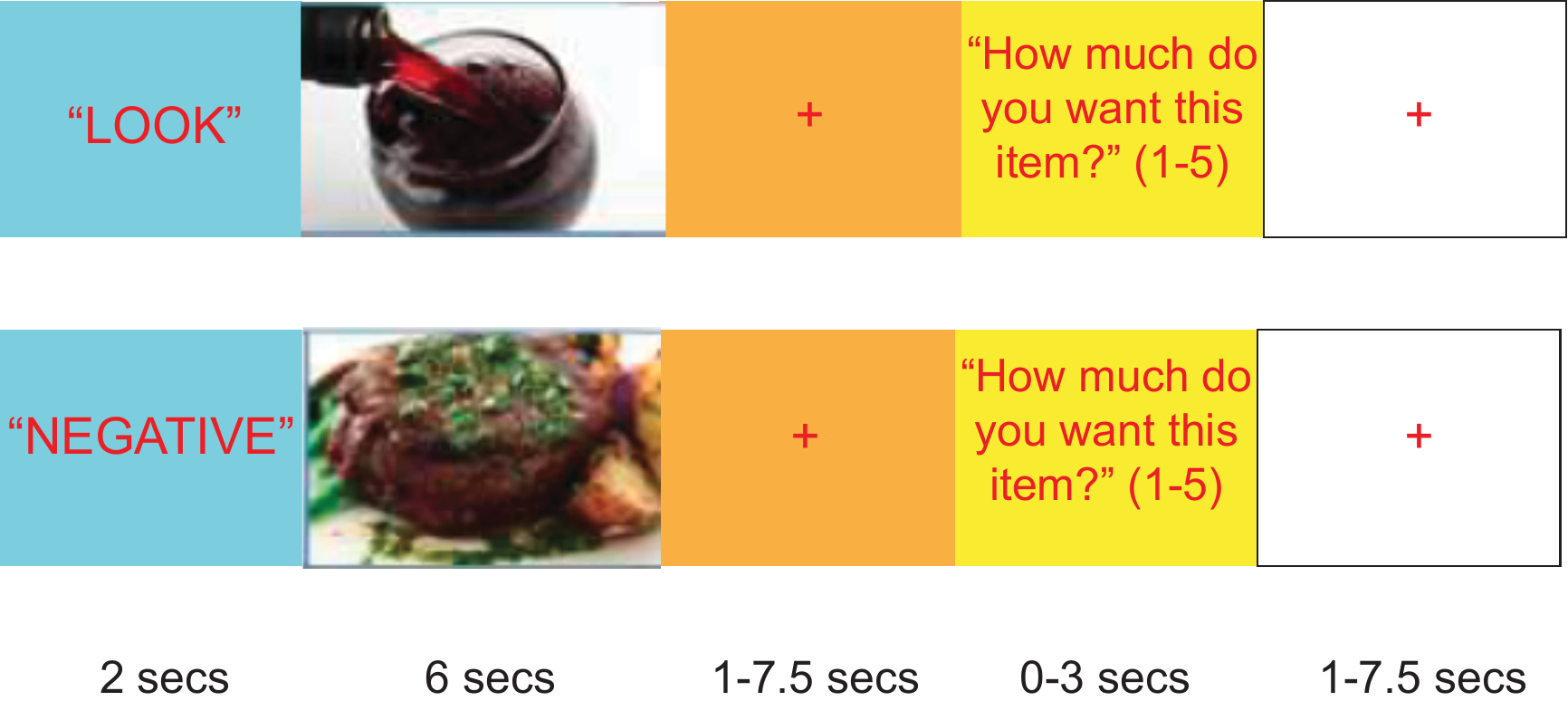

**Figure S2: CONSORT diagram of study**

### Analysis

Excluded (n=407)

♦  Not treatment seeking/enrolled in another trial (n=108)

♦  Did not complete screening process (n= 66)

♦  Did not meet minimum AUD criteria (n= 75)

♦  Excluded for psychiatric/medical conditions, cognitive impairment, or other exclusion criteria (n=158)

### Enrollment

Enrolled (n= 38)

Assessed for eligibility (n=445)

Did not undergo fMRI scanning (n= 3)

♦ Medical instability (n=1)

♦ Tattoo (n=1)

♦ Head too large for scanner (n=1)

Underwent baseline fMRI scanning (n=35)

Analyzed (n= 22)

♦ Excluded from analysis (dropped out of study) (n=12)

♦ Excluded from analysis (no post-CBT assessments) (n=1)

♦ Excluded from analysis (CIWA-AR>9 after 24 hours of abstinence (n=0)

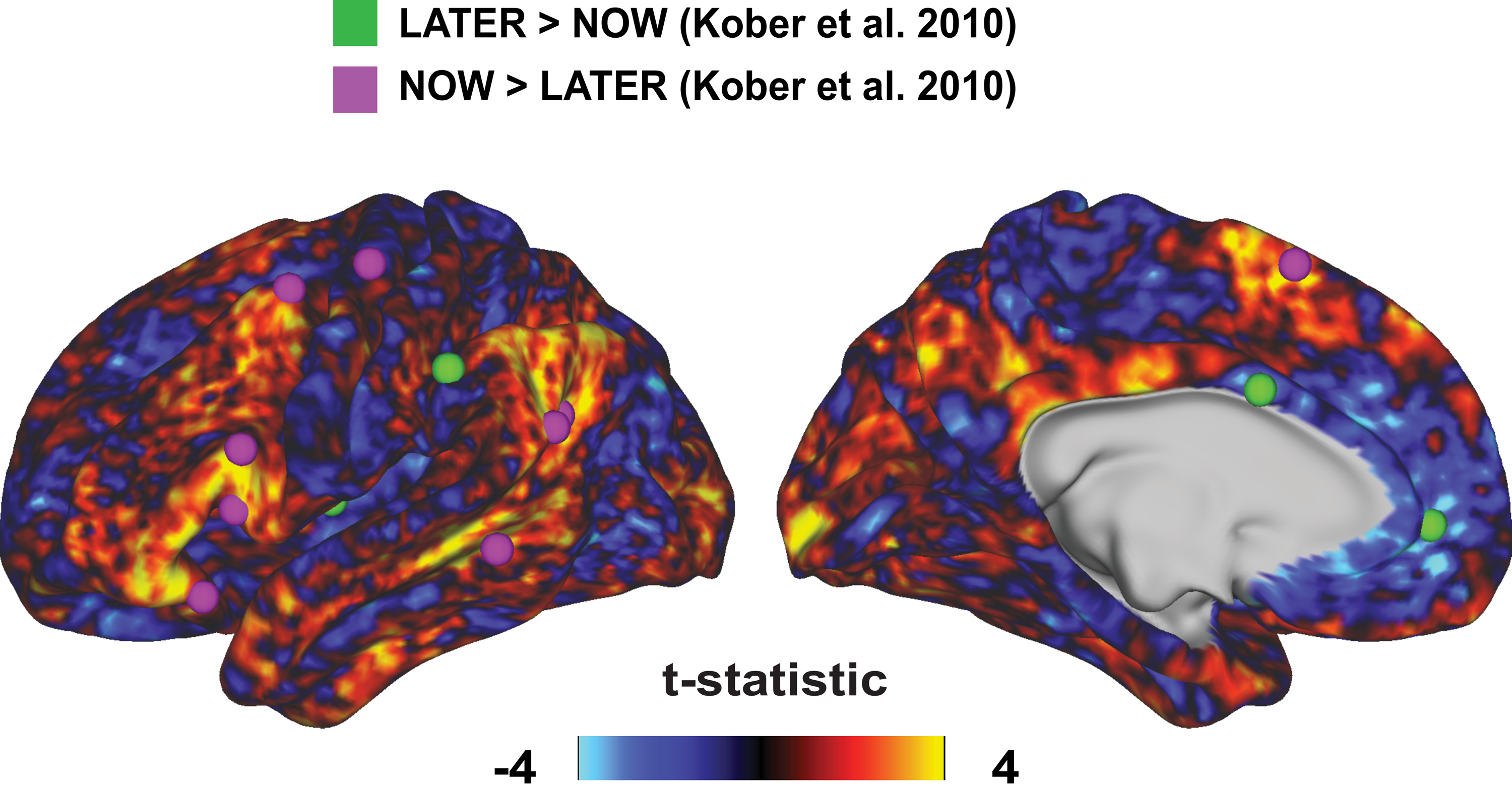

**Figure S3: T-statistical map of NEGATIVE-LOOK instructions.** Plotted are peak MNI coordinates from significant activations in the LATER-NOW contrast from Kober et al 2010 (corresponding to NEGATIVE-LOOK in our study).

| **ROI** | **F-stat** | | **P value (uncorr.)** | **P value (corr.)** | **Post-hoc finding** | **Post-hoc t-statistic** | **dF** | **Post-hoc p-value** |
| --- | --- | --- | --- | --- | --- | --- | --- | --- |
|  | | **Main effect of instruction** | | | | | | |
| L_STSvp | 22.175 | | 0.0001 | 0.0473 | NEGATIVE> LOOK | -5.45 | 87 | <0.0001 |
| **Main effect of cue type** | | | | |  |  |  |  |
| R_MST | 23.8316 | | 0.0001 | 0.0041 | Alcohol > Food | 5.9738 | 87 | <0.0001 |
| R_V2 | 9.5725 | | 0.0055 | 0.0447 | Food > Alcohol | -3.2166 | 87 | 0.0018 |
| R_V3 | 32.2525 | | <0.0001 | 0.0012 | Food > Alcohol | -6.0905 | 87 | <0.0001 |
| R_V4 | 19.7095 | | 0.0002 | 0.0055 | Food > Alcohol | -4.8169 | 87 | <0.0001 |
| R_V8 | 16.6244 | | 0.0005 | 0.0097 | Food > Alcohol | -5.3083 | 87 | <0.0001 |
| R_RSC | 11.3301 | | 0.0029 | 0.0289 | Alcohol > Food | 3.8440 | 87 | 0.0002 |
| R_POS2 | 12.8710 | | 0.0017 | 0.0191 | Alcohol > Food | 4.4029 | 87 | <0.0001 |
| R_IPS1 | 15.7389 | | 0.0007 | 0.0116 | Food > Alcohol | -4.5089 | 87 | <0.0001 |
| R_V3B | 13.1838 | | 0.0016 | 0.0188 | Food > Alcohol | -4.2186 | 87 | <0.0001 |
| R_MT | 14.0323 | | 0.0012 | 0.0157 | Alcohol > Food | 5.1668 | 87 | <0.0001 |
| R_7m | 24.0760 | | 0.0001 | 0.0041 | Alcohol > Food | 4.8472 | 87 | <0.0001 |
| R_v23ab | 23.3786 | | 0.0001 | 0.0041 | Alcohol > Food | 4.4346 | 87 | <0.0001 |
| R_6v | 9.1094 | | 0.0065 | 0.0480 | Food > Alcohol | -3.1891 | 87 | 0.0020 |
| R_a24 | 11.8733 | | 0.0024 | 0.0259 | Alcohol > Food | 3.1460 | 87 | 0.0023 |
| R_10r | 11.0495 | | 0.0032 | 0.0304 | Alcohol > Food | 3.6237 | 87 | 0.0005 |
| R_PoI2 | 21.0755 | | 0.0002 | 0.0047 | Food > Alcohol | -4.1388 | 87 | 0.0001 |
| R_PFt | 13.7447 | | 0.0013 | 0.0164 | Food > Alcohol | -3.8733 | 87 | 0.0002 |
| R_PH | 14.1324 | | 0.0012 | 0.0157 | Food > Alcohol | -3.5642 | 87 | 0.0006 |
| R_TPOJ2 | 22.6911 | | 0.0001 | 0.0042 | Alcohol > Food | 6.0336 | 87 | <0.0001 |
| R_TPOJ3 | 20.5977 | | 0.0002 | 0.0047 | Alcohol > Food | 5.5077 | 87 | <0.0001 |
| R_VMV3 | 14.8729 | | 0.0009 | 0.0139 | Food > Alcohol | -3.3403 | 87 | 0.0012 |
| R_V4t | 11.3650 | | 0.0029 | 0.0289 | Alcohol > Food | 3.1810 | 87 | 0.0020 |
| R_V3CD | 47.1333 | | <0.0001 | 0.0003 | Food > Alcohol | -8.3162 | 87 | <0.0001 |
| R_VM2 | 11.6969 | | 0.0026 | 0.0268 | Food > Alcohol | -3.7910 | 87 | 0.0003 |
| R_31pd | 12.9605 | | 0.0017 | 0.0190 | Alcohol > Food | 4.5096 | 87 | <0.0001 |
| R_31a | 10.1236 | | 0.0045 | 0.0378 | Alcohol > Food | 3.5192 | 87 | 0.0007 |
| R_25 | 10.8222 | | 0.0035 | 0.0322 | Alcohol > Food | 3.3851 | 87 | 0.0011 |
| L_V1 | 21.8443 | | 0.0001 | 0.0043 | Food > Alcohol | -4.5723 | 87 | <0.0001 |
| L_MST | 34.8102 | | <0.0001 | 0.0010 | Alcohol > Food | 6.0497 | 87 | <0.0001 |
| L_V2 | 16.9651 | | 0.0005 | 0.0092 | Food > Alcohol | -4.5342 | 87 | <0.0001 |
| L_V3 | 18.5159 | | 0.0003 | 0.0066 | Food > Alcohol | -5.3574 | 87 | <0.0001 |
| L_V4 | 39.9164 | | <0.0001 | 0.0006 | Food > Alcohol | -6.3051 | 87 | <0.0001 |
| L_V8 | 19.5580 | | 0.0002 | 0.0055 | Food > Alcohol | -5.3407 | 87 | <0.0001 |
| L_V3B | 14.1983 | | 0.0011 | 0.0157 | Food > Alcohol | -3.8941 | 87 | 0.0002 |
| L_MT | 22.0153 | | 0.0001 | 0.0043 | Alcohol > Food | 5.8492 | 87 | <0.0001 |
| L_7m | 20.6088 | | 0.0002 | 0.0047 | Alcohol > Food | 4.4005 | 87 | <0.0001 |
| L_v23ab | 15.5688 | | 0.0004 | 0.0082 | Alcohol > Food | 3.7665 | 87 | 0.0003 |
| L_d23ab | 9.3862 | | 0.0059 | 0.0458 | Alcohol > Food | 3.8600 | 87 | 0.0002 |
| L_31pv | 10.1645 | | 0.0044 | 0.0378 | Alcohol > Food | 3.7715 | 87 | 0.0003 |
| L_10r | 9.2460 | | 0.0062 | 0.0473 | Alcohol > Food | 3.3402 | 87 | 0.0012 |
| L_PFt | 13.7049 | | 0.0013 | 0.0164 | Food > Alcohol | -3.6567 | 87 | 0.0004 |
| L_TE1p | 9.4967 | | 0.0057 | 0.0448 | Alcohol > Food | 3.5632 | 87 | 0.0006 |
| L_TF | 11.1403 | | 0.0031 | 0.0302 | Alcohol > Food | 2.7859 | 87 | 0.0066 |
| L_TPOJ2 | 23.1638 | | 0.0001 | 0.0041 | Alcohol > Food | 5.7311 | 87 | <0.0001 |
| L_TPOJ3 | 27.6686 | | <0.0001 | 0.0026 | Alcohol > Food | 5.8468 | 87 | <0.0001 |
| L_PGi | 9.5560 | | 0.0055 | 0.0447 | Alcohol > Food | 3.6351 | 87 | 0.0005 |
| L_PGs | 10.3842 | | 0.0041 | 0.0367 | Alcohol > Food | 3.6012 | 87 | 0.0005 |
| L_VMV3 | 19.2895 | | 0.0003 | 0.0056 | Food > Alcohol | -4.7631 | 87 | <0.0001 |
| L_pOFC | 12.9780 | | 0.0017 | 0.0190 | Alcohol > Food | 4.3060 | 87 | <0.0001 |
| L_V3CD | 10.3248 | | 0.0042 | 0.0367 | Food > Alcohol | -3.7102 | 87 | 0.0004 |
| L_31pd | 14.5193 | | 0.0010 | 0.0150 | Alcohol > Food | 4.2274 | 87 | 0.0001 |
| L_25 | 15.9613 | | 0.0007 | 0.0113 | Alcohol > Food | 3.5517 | 87 | 0.0006 |
| L_s32 | 15.5048 | | 0.0008 | 0.0119 | Alcohol > Food | 4.1805 | 87 | 0.0001 |
| L_Ig | 9.1732 | | 0.0064 | 0.0477 | Food > Alcohol | -2.0770 | 87 | 0.0408 |

**Table S1: Parcels significant for main effects of instruction and cue type with post-hoc tests.**
